# Identification and characterization of the T cell receptor (TCR) repertoire of the Cynomolgus macaque (*Macaca Fascicularis*)

**DOI:** 10.1101/2022.03.30.486315

**Authors:** Swati Jaiswal, Shayla Boyce, Sarah K. Nyquist, Tasneem Jivanjee, Samira Ibrahim, Joshua D. Bromley, G. James Gatter, Hannah P. Gideon, Kush V. Patel, Sharie Keanne C. Ganchua, Bonnie Berger, Sarah M. Fortune, JoAnne L. Flynn, Alex K. Shalek, Samuel M. Behar

## Abstract

**Background:** Non-human primates (NHP) are desirable as animal models of human disease because they share behavioral, physiological, and genomic traits with people. Hence, NHP recapitulate manifestations of disease not observed in other animal species. The *Macaca fascicularis* (i.e., Cynomolgus macaque) is an NHP species extensively used for biomedical research, but the TCR repertoire hasn’t been characterized yet.

**Result:** We used the genomic sequences to design primers to identify the expressed TCR repertoire by single cell RNAseq. The data analysis from 22 unique samples were used to assign a functional status to each TCR genes. We identified and analyzed the TRA/D, TRB and TRG loci of the Cynomolgus macaque.

**Conclusion:** The genomic organization of the Cynomolgus macaque has great similarity with *Macaca mulatta* (i.e., Rhesus macaque) and they shared >90% sequence similarity with the human TCR repertoire. These data will facilitate the analysis of T cell immunity in Cynomolgus macaques.

## Background

Experimental animal models are an essential tool in our pursuit of understanding human physiology. The mouse has been incredibly useful in elucidating the major concepts of immunology, including defining the genetic and molecular basis of immunoglobulin and TCR formation and diversity. As part of this effort, the murine TCR repertoire have been extensively characterized and its knowledge is being used to develop new approaches to facilitate antigen discovery and novel treatments for human disease. However, it is not surprising that many human diseases are inadequately modelled in mice. This has been repeatedly emphasized for cancer and is also true for many infectious diseases. Two important examples are acquired immunodeficiency syndrome (AIDS), which is caused by the Human Immunodeficiency Virus-1 (HIV-1), and COVID-19, which is caused by the SARS-CoV2 coronavirus [1–5]. Mice are naturally resistant to both infections. For HIV research, the field largely turned to nonhuman primates (NHP) as a better alternative because they could be infected with a highly related virus, Simian Immunodeficiency Virus (SIV). Consequently, the Rhesus macaque’s TCR locus was among the first NHP TCR locus to be characterized [6]. Cynomolgus macaques have been increasingly used for biomedical research, especially in the fields of neurology, cardiology, and for drug development [7, 8]. Importantly, they are increasingly used for infectious disease research, including as a model for human HIV [9] and SARS-CoV2 infection [5]. Most NHP species, including rhesus macaques, whether in captivity or in the wild, rapidly succumb to *Mtb* infection [10, 11]. However, Flynn’s group finds that following challenge with *Mtb*, 50% of infected Cynomolgus macaques develop a form of disease that resembles latent TB in people [12–15]. Indeed, the pathology observed among *Mtb*-infected Cynomolgus macaques *recapitulates* the entire spectrum of pathology of human TB granulomas [16]. Thus, the Cynomolgus macaque is providing insights into human disease not possible with other small animal models.

The tremendous capacity of T cells to recognize diverse antigens has a genetic basis that is inherent in the genomic organization of the T cell receptor (TCR) loci [17]. TCR repertoire diversity arises through genetic mechanisms that minimize the number of genetic elements encoded by the genome while maximizing the potential breadth of expressed TCRs. The germline configuration of TCR genes is not functional. Instead, the TCR loci encode families of variable (V), diversity (D), and joining (J) segments, which undergo rearrangement early during T cell development [17]. Recombination of V, D, and J segments leads to a gene fragment that encodes the V-region domain, which becomes the N-terminus of the TCR protein and determines its antigen specificity. Downstream of the V, D, and J genes are constant (C) region exons, which encode the C-terminus of all TCRs and couples the TCR to the Cluster of differentiation 3 (CD3) protein complex to mediate signal transduction into the cell. The primary diversity of TCRs arises from the nearly random assortment of V, D, and J gene segments, as well as additional diversity that occurs at the V-D and D-J junctions by imprecise recombination and the insertion of non-germline encoded nucleotides (N-regions). TCRs are heterodimers formed by TCRα and TCRβ chains, which are encoded by distinct loci (TRA and TRB, respectively) [18]. The TCRα is encoded when Vα and Jα gene segments recombine; the TCRβ is formed from the recombination of Vβ, D and Jβ gene segments. Additional diversity is created by the random pairing of the TCRα and TCRβ chains. Unlike immunoglobulin genes, somatic mutation does not occur in TCR genes. The potential TCR repertoire varies between animal species and is driven in large part by the number of functional members of V, D, or J segments. In humans, there is the potential to generate 10^15^ unique TCRs.

A second subset of T cells are known as Gamma-delta (γδ) T cells, express an alternative TCR, which is encoded by distinct gene segments found in the TRG and TRD loci. The γδ-TCR is structurally similar to the αβ-TCR. Like the TRA and TRB loci, the TRG and TRD loci contain sets of Vγ and Jγ, and Vδ, Dδ and Jδ gene segments, respectively. In general, there are fewer gene segments in the TRG and TRD loci [19]. γδ T cells remain enigmatic because the antigens they recognize and the antigen presenting molecules that restrict their recognition of antigen are incompletely characterized. Nevertheless, they are identified in the circulation and in the tissues of all mammals, and play important roles in autoimmune disease, and in immunity to infection and cancer [20, 21].

Here we identified the TRA, TRB, TRG and TRD loci of the Cynomolgus macaque. Based initially on the homology with human TCR gene segments, and subsequently using the identified gene segments from Rhesus macaque and Cynomolgus macaque, we systematically identified all the V, D, J, and C gene segments belonging to all four T cell receptor loci. Finally, using the genomic sequences, we designed specific primers for the amplification of the Vα and Vβ regions, and determined which of the V gene segments are expressed in individual subjects. These data will allow the detailed analysis of the T cell responses in Cynomolgus macaques as well as comparative immunogenetics studies, comparing different species of Cynomolgus macaques, as well as the evolution of TCR genes among the primates.

## Results

### Identification of the *<u>Mac</u>aca <u>fas</u>cicularis* (macfas) TCR loci

Based on nucleic acid sequence homology with the human Cα, Cδ, Cβ, and Cγ gene segments, the TRA and TRD loci were identified on Chr.7, and TRB and TRG loci were identified on Chr.3 (Figure 1). Subsequently, each human V, D, J, and C gene segment was used to blast the macfas Chr.7 and 3, to identify homologous gene segments. Similarly, *<u>Mac</u>aca <u>mul</u>atta* (macmul) gene segments were also used to identify homologous genes unique to the macaca genus. Using this approach, we were able to annotate and assemble a map of the macfas TRA, TRB, TRG, and TRD loci as described in detail below.

**Figure 1:**
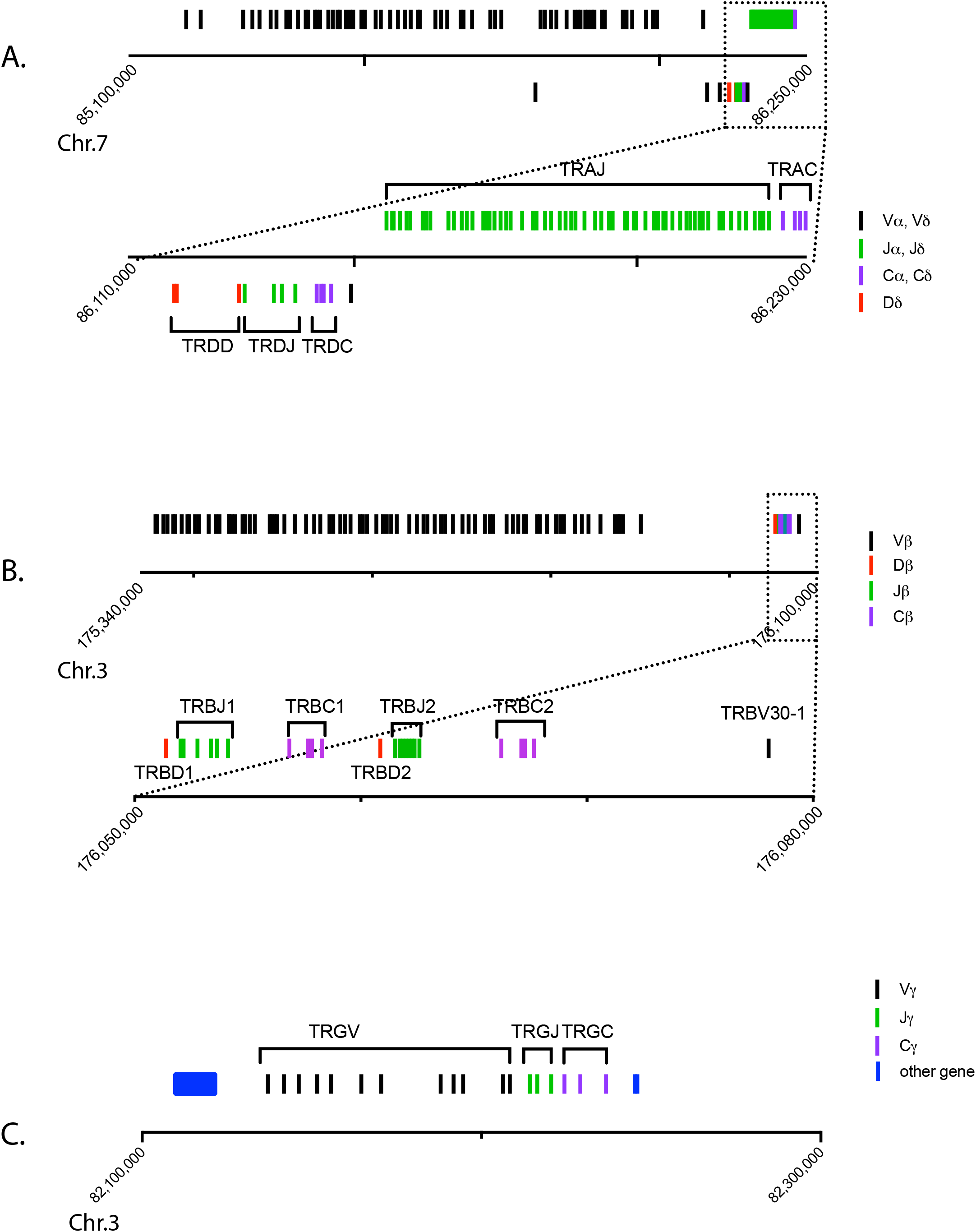
The Macfas TCR loci. Structure of the TCR loci (A) TRA/TRD (B) TRB and (C) TRG loci. (A) The TRA and TRD loci are interspersed on Chr. 7. The genes above the x-axis belong to the TRA locus; those below the axis belong to the TRD locus. The boxed region is expanded to show greater detail. (B) The TRB locus is located on Chr. 3. The boxed region is expanded to show greater detail. (C) The TRG locus is located on Chr. 3. Each line represents a gene and the distance between them is proportional to their spacing on Chr.7 and Chr.3. The blue boxes represent the 3′ region of the AMPH gene and exon 10 of STARD3NL, which are boundaries of the TRG locus. The black lines represent V gene, green lines are J chain, purple lines represent C region, and the red lines are representing D region.

### The macfas TRA locus

The structure of the macfas TRA locus is like the human locus in that it overlaps the TRD locus on Chromosome 7 (Figure 1A) [22]. We identified 62 TRAV genes in macfas, one more than the 61 human genes but less than the 67 macmul genes. The two human gene families TRAV7 and TRAV28, each contain a single member and are absent from the macfas and macmul TRA locus (Table 1, Figure 2). Conversely, the TRA loci of macfas and macmul have one additional gene in the TRAV24, TRAV25, and TRAV26 families. The greater number of macmul TRAV genes compared to macfas results from an expansion of the TRAV22 and TRAV23 families, from one member to three and four, respectively (Table 1). Of the 62 macfas TRAV genes, 12 are pseudogenes and 2 are ORFs (Table 1, Table S1). There might have been a gene duplication of the TRAV genes TRAV24, TRAV25, TRAV26, which differentiates the human TRAV locus from the macaque locus (Figure 2B). Second, there are additional members of the TRDV1. TRAV22, and TRAV23 families in NHP, compared to the human TRA/DV locus. We did not find macfas homologs of macmul TRDV1-1, TRAV22-2, TRAV22-3, TRAV-23-2, TRAV23-3, or TRAV23-4, despite searching the macfas genome using the macmul homologs. We believe the differences between the macfas and macmul genomic sequences could have arisen from gaps in the known macfas sequence or problems with the genome assembly.

**Table 1:**
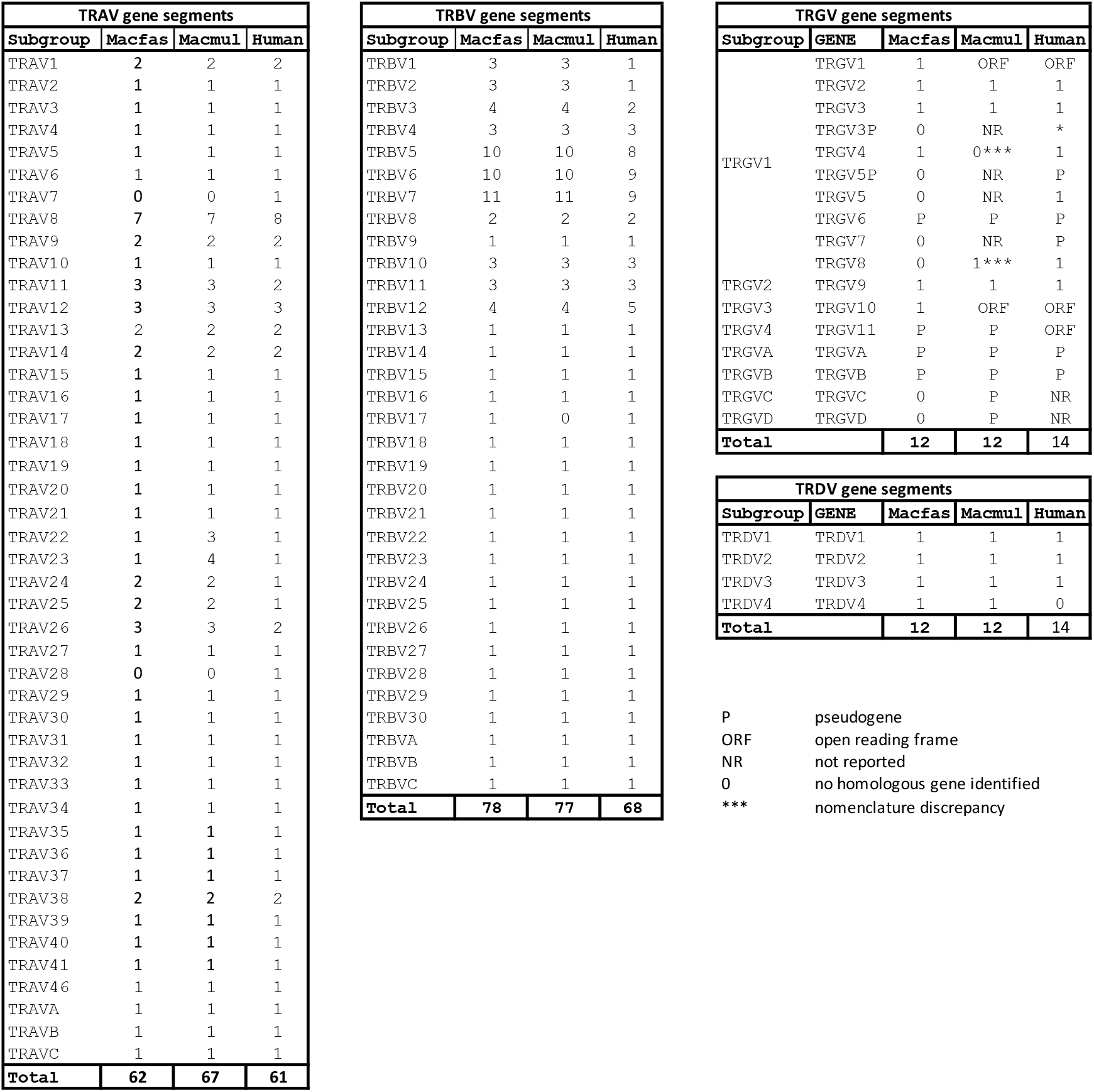
Comparison of genes (TRAV/ TRBV/ TRDV/ TRGV) in macfas, macmul and human

**Figure 2.**
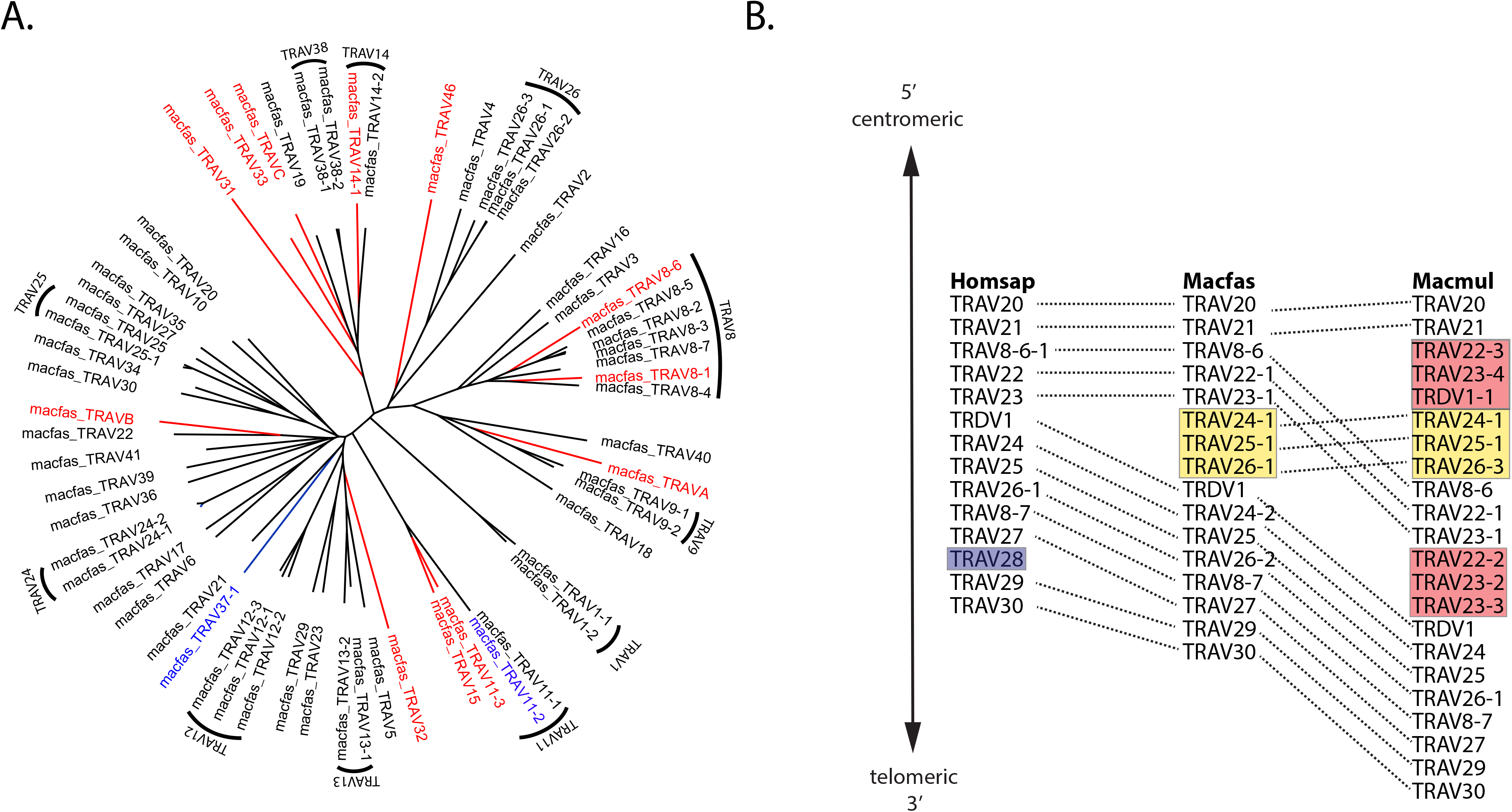
TRAV families. Phylogenetic tree illustrating (A) functional genes (black), pseudogenes (red) and ORFs (blue) of the macfas TRAV locus. The genes clustered together belong to the same family. (B) Comparison of the TRAV/TRDV locus of homsap, macfas, and macmul. The genes that are exclusive to humans are highlighted in purple. Those TRAV genes found in macfas and macmul but not in homsap are in yellow, and the genes are present only in macmul but absent in macmul are in red. See text for the details.

We identified 61 TRAJ genes, which is the same number as Rhesus macaque and human TRAJ genes. There is a high degree of conservation between macfas and *<u>Hom</u>o <u>sap</u>iens* (homsap) TRAJ gene segments (Table S2). Finally, we compared the TRAC exons from all three species. The macfas and macmul TRAC genes have identical amino acid sequences (Figure S1).

### The macfas TRB locus

The macfas TRB locus (Figure 1B) is similar in structure to the macmul TRB locus. We identified 78 TRBV genes, compared to 77 annotated macmul TRBV genes (Table 1 & Table S3). Both are expanded compared to the human species, for which there exists 68 distinct genes. The overall TRBV family structure is similar, with some variation in the number of members and the number of pseudogenes (n=17) and ORFs (n=2) (Table 1, Figure 3). The organization of the TRBJ and TRBC genes is similar in all three species, characterized by a duplication of the TRBJ and TRBC genes (Figure 1B). Comparing the macfas and macmul TRBJ gene segments, four (including the TRBJ2.2P ORF) differ by a single nucleotide; the other 10 genes are 100% conserved (Figure S2, Table S4). The TRBD1 and TRBD2 are also 100% conserved between macfas and macmul (Table S4). Similarly, there is a high degree of conservation between macfas and homsap TRBJ gene segments (Figure S2). Finally, we compared the TRBC exons from all three species. As noted, there are two TRBC genes, TRBC1 and TRBC2, which are 97% identical. The macfas and macmul TRBC1 differ by only two bp and the translated sequence is 100% identical; for TRBC2, there is a single aa difference (Figure S1).

**Figure 3.**
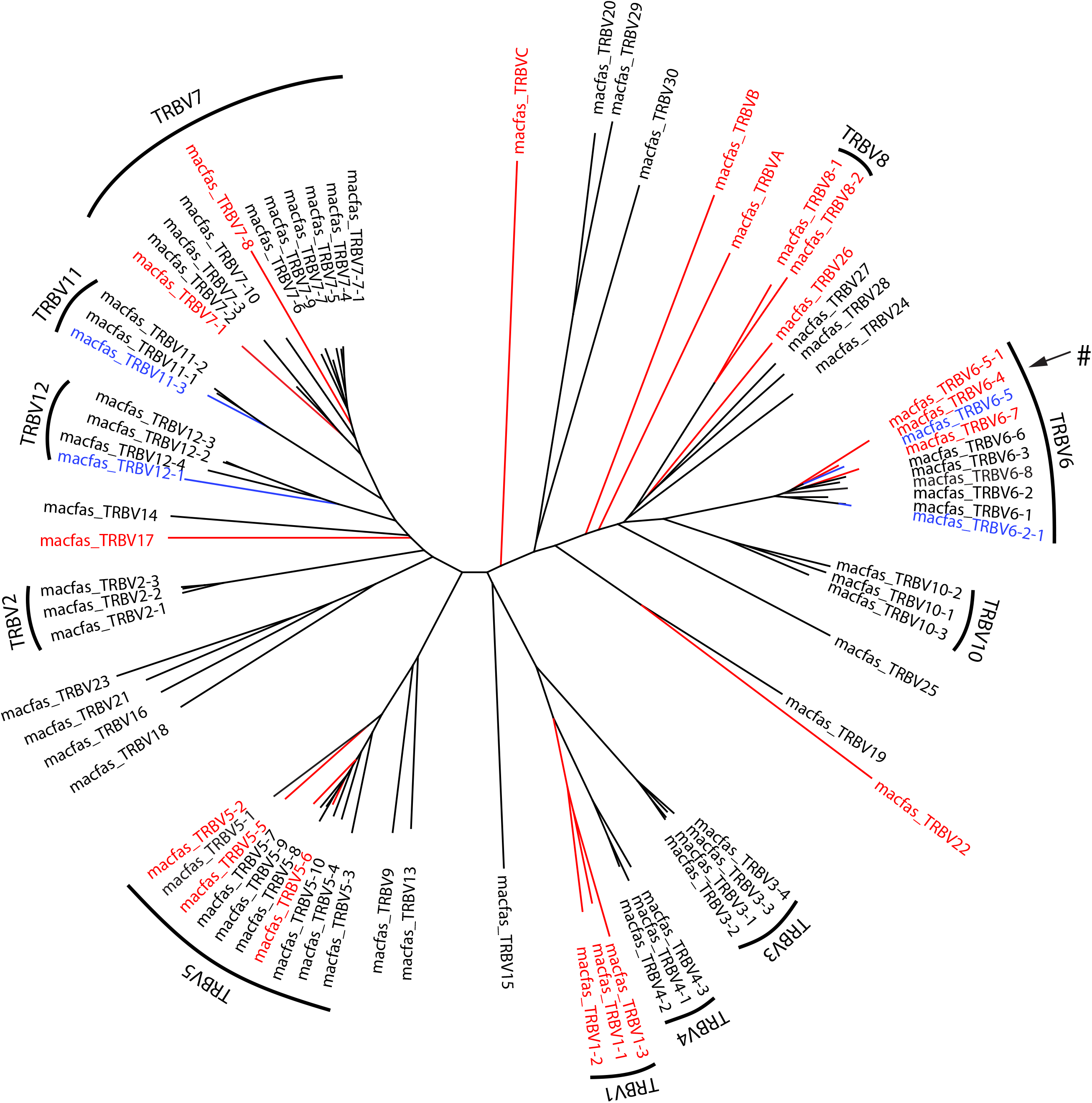
TRBV families. Phylogenetic tree illustrating functional genes (black), pseudogenes (red) and ORFs (blue) of the macfas TRBV locus. The genes clustered together belongs to the same family. #, TRBV6-4 is a pseudogene in the genomic sequence, but the expressed gene is functional (see results, “Expressed V gene repertoire”).

### The macfas TRG locus

The macfas TRG locus is located on chromosome 3 (Figure 1C). We identified 12 TRGV genes of which 6 are predicted to be functional and an additional 4 are pseudogenes (Figure 4, Table 1, Table S5). These genes were compared to the homologous genes in human and rhesus (Figure 4). The same 12 genes were found in the macmul TRG locus. We named macfas TRGV4 based on its homology with homsap TRGV4*01 (92% homology, Figure 4). The ortholog in the macmul TRG locus is identical in sequence, but IMGT names it TRGV8, although it has only 88% homology to homsap TRGV8*01. In general, the macfas and macmul orthologs had between 0-2 mismatches (i.e., >99% homology), while the homology between macfas and homsap TRGV genes was 88-95%. The two NHP species lacked TRGV5, TRGV5P, TRGV7, and TRGV8, and macmul had two additional V genes, TRGVC and TRGVD. The human TRG locus has two clusters of J segments and C-region genes [22, 23]; IMGT/LIGM-DB: IMGT000011 (582960 bp), human (Homo sapiens) TRG locus), and the macmul locus has a similar structure (IMGT/LIGM-DB: IMGT000059 (197016 bp), Rhesus monkey (Macaca mulatta) TRG locus). While there are five macmul TRGJ gene segments, we detected only three in the macfas TRG locus. These three are more like the macmul TRGJ2-1, 2-2, and 2-3 gene segments (Figure 4B). Similarly, there is a single macfas TRGC region gene, and its amino acid sequence is 91.9% and 96.5% identical to macmul TRGC1 and TRGC2, respectively (Figure S1). There is 1, 0 and 2 mismatches between macfas and macmul TRGC2 exon 1, 2, and 3, respectively. The human TRGC2 exon 2 contains a duplicated sequence, which neither the macfas nor macmul exon 2 contains (Figure S2). Therefore, the macfas TRGC gene is the TRGC2 ortholog, and the genomic assembly of macfas is missing a region that spans TRGJ1-1 to TRGC1 (Figure 4C).

**Figure 4.**
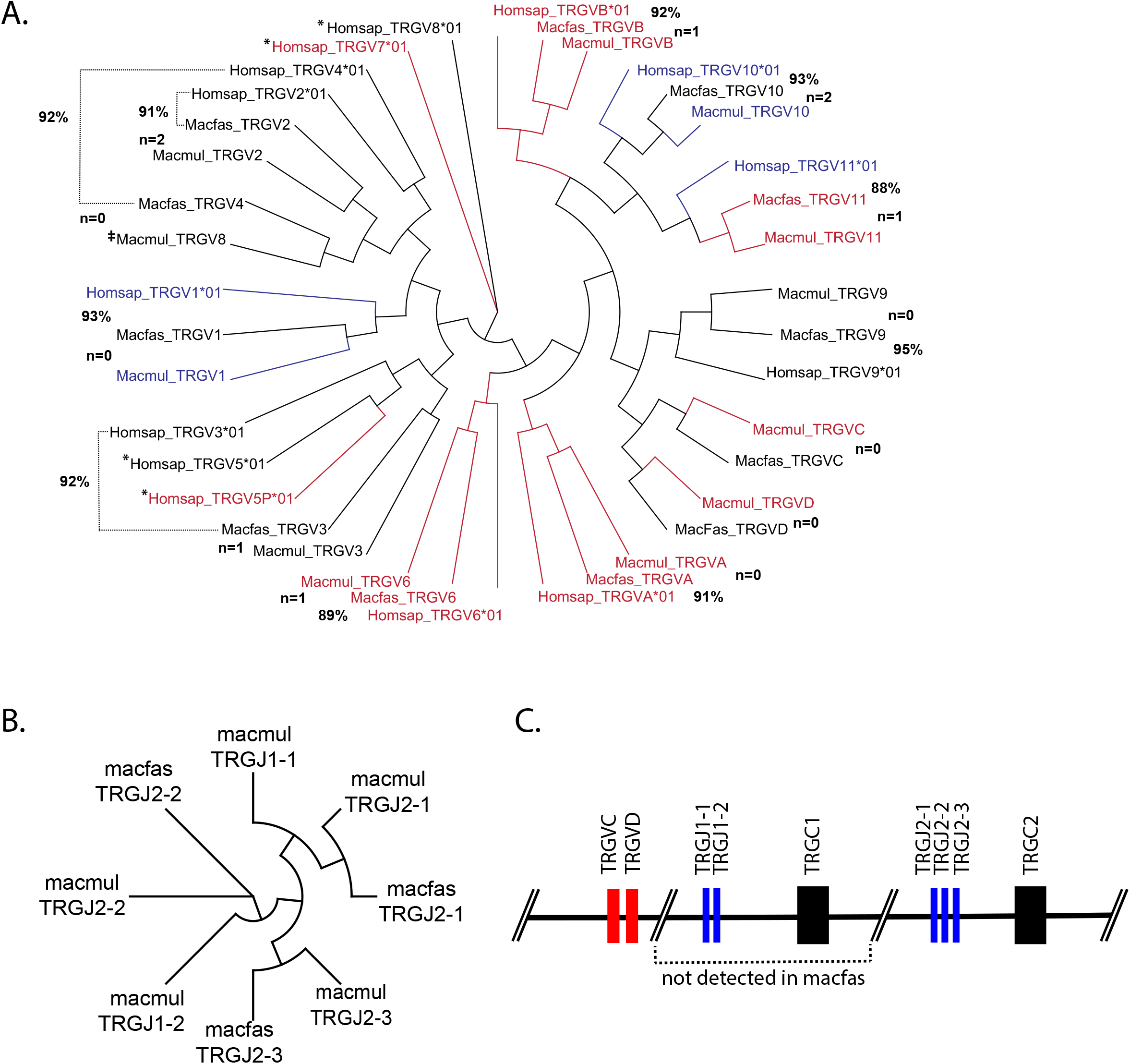
TRGV and TRGJ gene segment homologies. (A) Phylogenetic tree illustrating functional genes (black), pseudogenes (red) and ORFs (blue) of the macfas TRGV locus. The number (i.e., “n=1”) is the number of mismatches between the macfas and macmul genes. The % is the homology between the macfas and the homsap gene. Homologies between other genes of interest are indicated with a dotted line. *, homsap genes for which no macfas or macmul homologs were identified. ‡, nomenclature discrepancy. (B) Phylogenetic tree clustering macfas and macmul TRGJ genes. (C) Schematic of the genomic organization of the 3’ region of the TRG locus. TRGV (red), TRGJ (blue) and TRGC (black).

### The macfas TRD locus

The macfas TRD locus is located on chromosome 7 and overlaps with the TRA locus (Figure 1A). Three canonical TRDV genes were identified as macfas homologs of homsap TRDV1, TRDV2, and TRDV3, with homologies between 91-97% (Figure 5, Table S6). A fourth gene, TRDV4, was identified which was 100% homologous to macmul TRDV4. The macmul genome has a fifth gene, TRDV1-1, which is very homologous to TRDV1 (Figure 2, 5); no macfas orthologs were found for this gene. Three TRDD and four TRDJ macfas gene segments were identified, as in the homsap genome (Table S6). These genes are 100% identical to their macmul homologs. Similarly, the single macfas TRDC region has 100% DNA sequence identity and predicted amino acid sequence as the macmul TRDC (Figure S1). There is a two amino acid gap, which we suggest is a consequence of the artificial splicing between exons 2 and exon 3.

**Figure 5.**
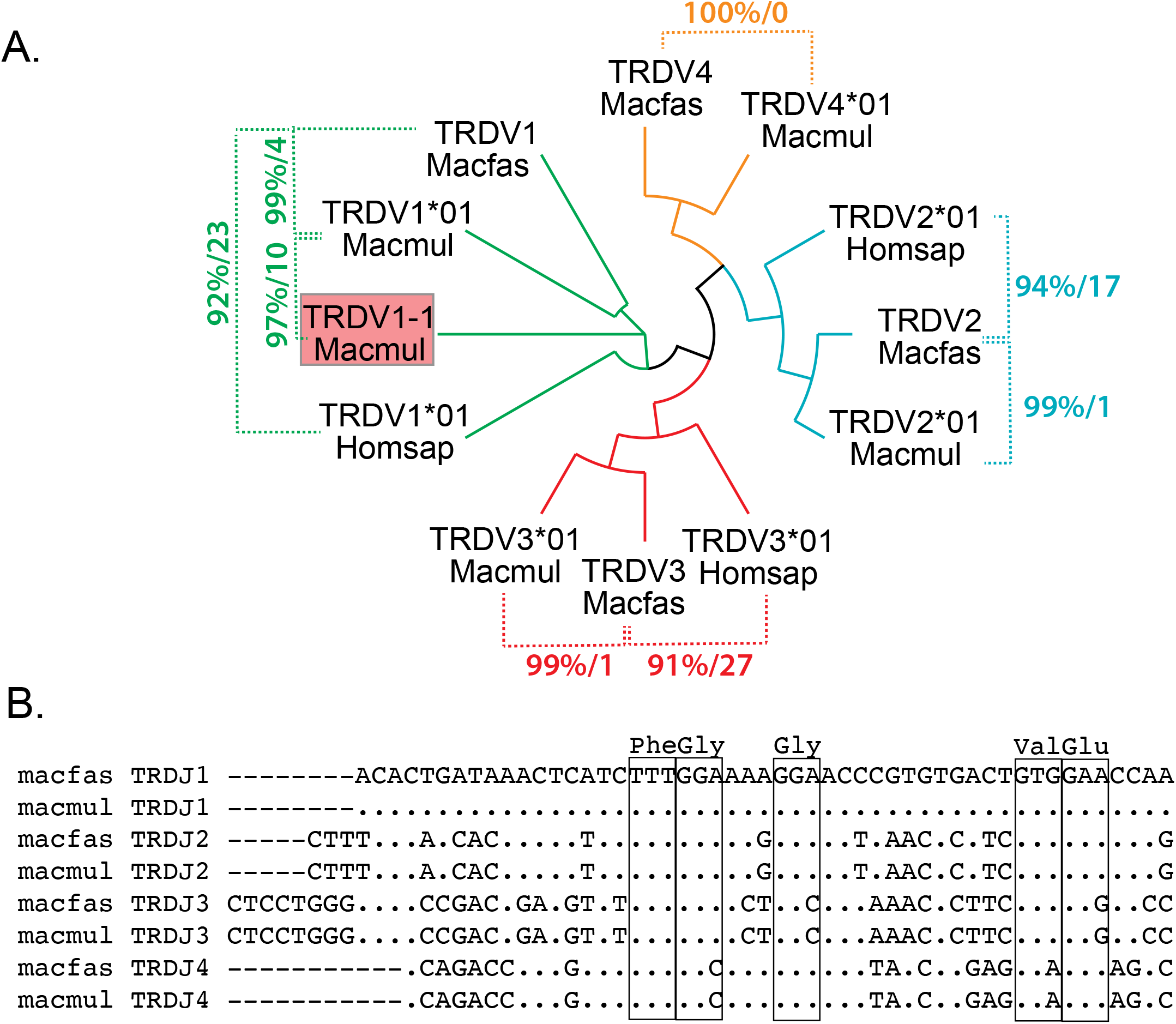
TRDV and TRDJ gene segment homologies. (A) Phylogenetic tree showing the functional genes homsap, macfas and macmul TRDV genes. Comparisons are indicated with dotted lines and the percent homology is indicated followed by the number of sequence mismatches. Each TRDV gene family is color coded. (B) Alignment of macmul and macfas TRDJ showing the conserved amino acids (boxed).

### The expressed V gene repertoire used by Cynomolgus macaque T cells

To determine the functionality of the TRAV and TRBV gene segments we identified, the following criteria were used: (i) Defined L1 exon and L2-V exon, (ii) absence of nonsense or missense mutation, and (iii) encodes a cytosine (C) at position 21-23 followed by tryptophan (W) at position 31-33 of the 3’ end. The terminal amino acids encoded by a functional TRAV gene is usually CAVR, CAL, or CAF. Similarly, the terminal amino acids encoded by a functional TRBV gene is usually CASSQ, CASSL, or CASSE. Based on these criteria, we initially assigned each V gene to be functional if it met these criteria. If the gene had an internal stop codon, or lacked the conserved C or W residue, it was deemed a pseudogene. Finally, if the gene appeared to be functional, but the L1 or L2 parts of the leader sequence could not be identified, or it lacked consensus splice site for intron A, we designated it an open reading frame (Supplemental tables 1, 3, and 5).

To determine the expressed TRAV and TRBV repertoire, TCRs from Cynomolgus macaques infected with *Mycobacterium tuberculosis* were analyzed. The distribution of the expressed TRAV and TRBV genes in uninvolved lung from these infected subjects was determined for 22 unique individuals (Figure 6). The distribution of TRAV and TRBV genes was also determined in BAL cells, before or after infection, involved (i.e., granulomas) lung tissue, and single cell lung suspensions (Figure S3). The presence of stop codons in pseudogenes that were observed to be transcribed were confirmed (data not shown). One exception was detected. For TRBV6-4, the expected stop codon at position 85 (TAG) was CAG in the transcribed gene, and thus, encoded a functional glutamine (Q). This difference between the germline and the transcribed gene could be the result of a polymorphism or a sequencing error in the genomic reference sequence. Finally, the status of V genes designated as ORFs, was changed to ‘functional’ if the V gene was transcribed.

**Figure 6.**
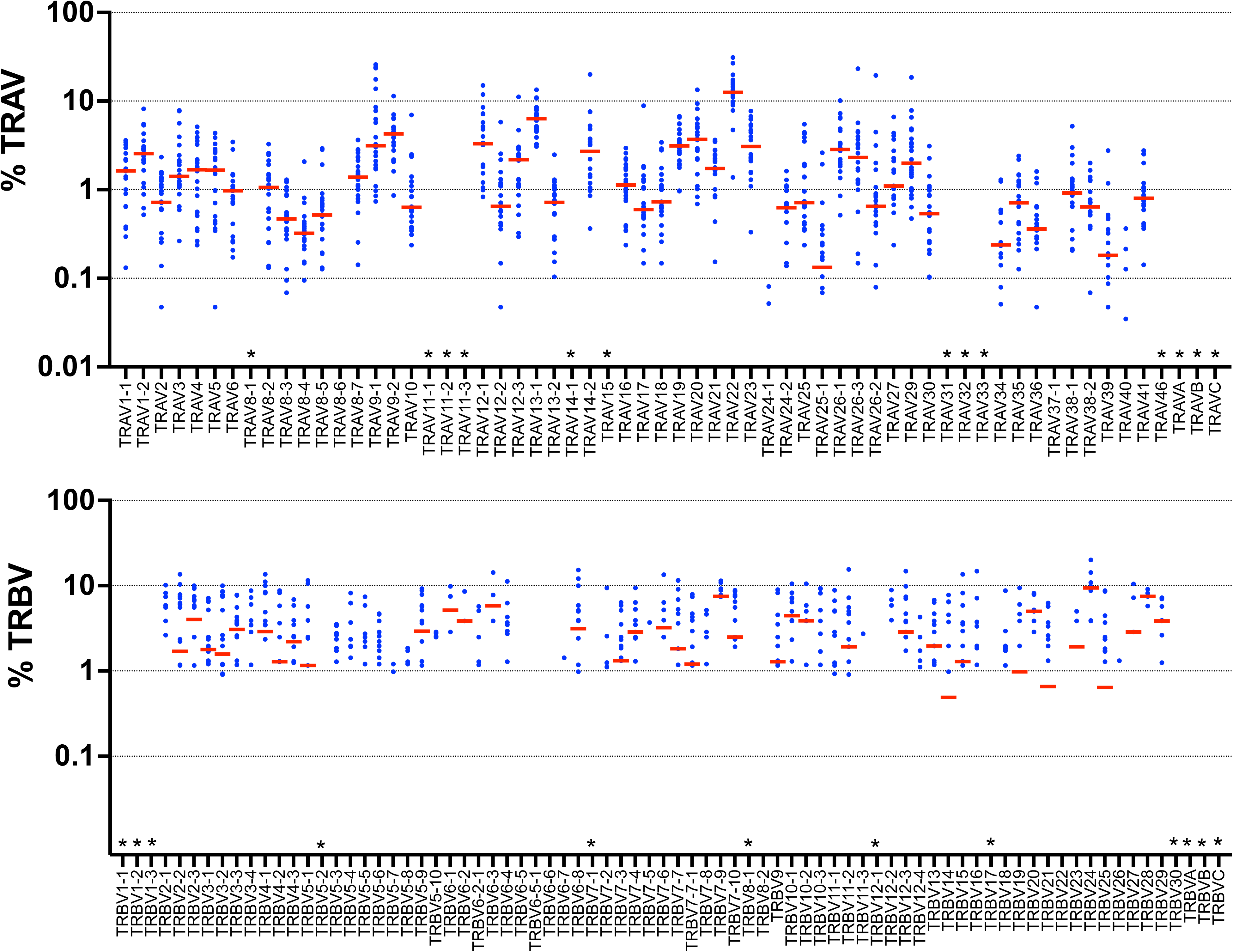
The expressed TRAV and TRBV repertoire. Single cell analysis of lung mononuclear cells from Cynomolgus macaques reveals their functionally expressed TRAV and TRBV repertoire. Each dot represents a different animal. All samples are from uninvolved lung tissue from subjects infected with *M. tuberculosis*. The average percentage was calculated for the TRAV (A) and TRBV (B) and the distribution was individually normalized for each subject. Red bar represents the median. *, expression not analyzed.

## Discussion

The nucleic acid sequence of recombined V, D, and J gene segments encodes the protein structure of the TCR and contains immunological information about T cell responses. The complementarity determining region 3 (CDR3), defined as the V-D-J or V-J recombination site, is unique to each unique T cell clone, sometimes referred to as a clonotype. Analytical approaches are beginning to predict the antigen specificity based on the primary sequence of the TCR. In the absence of the antigen specificity, the TCR sequence can be used as a surrogate of antigen specificity. As T cells undergo clonal expansion after encountering antigens, TCR sequences are being used to track T cells, monitor immune responses, and identify new antigens for human tumors and pathogens [24–26]. Advances in T cell therapy are being driven by our ability to clone and recombinantly express TCRs, as exemplified by adoptive cell therapy (ACT) [27, 28]. Thus, defining the V, D, and J gene segments is an important step in the analysis of T cell immunity.

We identified and annotated the TRA, TRB, TRG, and TRD loci of the Cynomolgus macaque. There is generally more than 90% homology between the different V, D, and J gene segments in the Human, Rhesus and Cynomolgus macaque’s TCR repertoire. As one might expect, the structure of the different TCR loci is highly conserved between Rhesus and Cynomolgus macaque. The differences we detected (e.g., Fig.2A) are more likely to be due to ascertainment bias arising from problems with genomic sequencing and assembly, than true evolutionary events. In support of this conclusion, among the expressed TCR repertoire, we found many macfas TCRs expressed in the lung matched to macmul reference sequences that were missing from the macfas genomic sequence. We also find that there is expansion of TCR beta locus of macfas and macmul compared to homsap. These differences, which are likely to have occurred by gene duplication [29, 30], may have occurred in response to changes in selective pressure during evolution of the TCR loci [31, 32].

## Conclusions

We identified and annotated the TRA/D, TRB and TRG loci of the Cynomolgus macaque. The TRA and TRB genomic sequences were used to design primers, and as reference sequences, to amplify and identify TCR sequences expressed by single cells from the lungs of Cynomolgus macaques. By using these data to analyze the αβTCRs expressed by mature T cells, we were able to discern which V genes were functional based on their RNA expression. This allowed us to refine and validate our predictions based on the genomic sequences. Altogether, these data show the utility of these TCR reference sequences, and we expect that they will be useful for the study of T cell immunity in Cynomolgus macaques.

## Methods

### Source of genomic sequence

The genome of the Cynomolgus macaque (NCBI: taxid 9541), also known as the crab eating macaque, has been sequenced and we used the reference genome Macaca_fascicularis_5.0 available from NCBI (https://www.ncbi.nlm.nih.gov/genome/776) [33, 34]. The formal genus and species name is *Macaca fascicularis*, which we abbreviate as macfas. The Rhesus macaque (i.e., *Macaca mulatta*; macmul) TCR sequences were obtained initially from the literature [35] and later from IMGT (http://www.imgt.org) [36]. The Human (i.e., *Homo sapiens*; homsap) TCR sequences were obtained from IMGT. In cases where more than one allele was available, the first allele was used for sequence comparisons.

### Annotation and analysis of Macfas TCR repertoire

To identify the location of the macfas TCR loci, the human TRAC, TRBC, TRGC, and TRDC were blasted against the macfas genome. Subsequently, all human gene segments were individually blasted against the macfas genome. As macfas gene segments were defined, they were also used to look for other homologous genes. At the beginning of this study, the sequences of the macmul TCBV genes were available and were used to look for homologous genes [35]. The names of the genes were assigned based on the homology with the human genes, and the location in the genome. The leader sequence (L1 & L2), TRV region, D region and J chain were identified for each gene. The annotation was done by following standard IMGT rules (http://www.imgt.org). Clustal Omega was used for multiple sequence alignments (https://www.ebi.ac.uk/Tools/msa/clustalo/) [37] and visualized using Archaeopteryx for Figures 2–5 [38]. Sequences were entered and tracked in SnapGene (version 5.0)

### Expressed TCR repertoire of Cynomolgus macaques

Cells from bronchoalveolar lavage (BAL), single cell suspensions of lung, or lung tissue, were obtained from Cynomolgus macaques infected with *Mycobacterium tuberculosis* and single cell RNAseq libraries were created [39]. Primers were synthesized that were specific for the different TRAV, TRBV, TRAC, and TRBC gene segments based on the genomic sequences described herein and used to enrich and amplify the TCR sequences from T cells in scRNA-Seq libraries generated using 3’ barcoded Seq-Well [40, 41]. Primers were not designed for pseudogenes that had internal stop codons, or for some V genes that were not initially identified. The libraries were sequenced and then aligned to the TCR reference sequences. The samples were analyzed for 48 TRAV and 73 TRBV genes. The V region and J region sequences were mapped using BOWTIE 2 as part of the TCRGO algorithm (https://github.com/ShalekLab/tcrgo/tree/master/tcrgo)[41]. Briefly, reads are aligned with the V and J regions in the reference TCR database, containing the sequences annotated in this report (see Results, below). Each read from a Seq-Well library includes nucleic acid tags that identify the cell of origin (cell barcode) and the transcript of origin (unique molecular identifier, UMI). Reads with matching cell barcode and UMI are merged, and a consensus V and J region mapping is determined based on sequence similarity identified among the majority of reads. A consensus CDR3 sequence is identified from reads with shared mappings.

## List of abbreviations

Adoptive cell therapy: (ACT)
Acquired immunodeficiency syndrome: (AIDS)
Constant: (C)
Cluster of differentiation 3: (CD3)
Complementary determining region 3: (CDR3)
Diversity: (D)
Human Immunodeficiency Virus-1: (HIV-1)
Joining: (J)
Gamma-delta: (γδ)
*Macaca fascicularis*: (macfas)
*Macaca mulatta*: (macmul)
*Mycobacterium tuberculosis*: (*Mtb*)
Non-human primates: (NHP)
Simian Immunodeficiency Virus: (SIV)
T cell receptor: (TCR)
Variable: (V)

## Declarations

### Ethics approval

All experiments, protocols, and care of animals were approved by the University of Pittsburgh School of Medicine Institutional Animal Care and Use Committee (IACUC). The Division of Laboratory Animal Resources and IACUC adheres to national guidelines established by the Animal Welfare Act (7 U.S. Code Sections 2131-2159) and the Guide for the Care and use of Laboratory Animals (Eighth Edition) as mandated by the U.S. Public Health Service Policy. The DLAR program is AAALAC accredited. Animals used in this study were housed in rooms with autonomously controlled temperature and provided enhanced enrichment procedures.

### Consent for publication

Not applicable

### Availability of data and materials

All data generated or analyzed during this study are included in this published article [and its supplementary information files]

## Authors’ contributions

S.J. and S.M.B., conceptualized and wrote the manuscript; S.B., helps in preliminary data acquisition; S.K.N., contributed to methodology and performed sc RNA seq data analysis; S.K.G., K.P., H.G. NHP Sample processing and facilitated transfer of the samples; T.J., S.I., J.B., and G.J.G., have contributed in T cell processing for scRNA sequencing; A.K.S. and B.B., provided resource and supervision for scRNA sequencing and data analysis; Funding Acquisition, J.F., S.M.F., S.M.B.; S.J. and S.M.B. have reviewed and edited the manuscript. All authors reviewed the manuscript.

## Acknowledgements

Roisin Floyd, Marc Wadsworth, and Travis Hughes have performed original sequence libraries for depletion experiment that were helpful in analyzing the expressed repertoire, Jake Rosenburg and Andy Tu helped in the development of TCR pipeline and interpretation of results.

## Notes

1. The numerical value for every gene represents the number of allele present.
2. ORF: Open reading frame
3. NR: Not reported
4. P: Pseudogene
5. F: Functional
6. ***: Nomenclature discrepancy

## Supplemental information

**Figure S1.** Constant region homology. Alignment of the amino acid sequence of the TCR constant regions, derived from the in silico splicing of the human, macfas and macmul TRAC, TRBC, TRGC, and TRDC exons. Dots represent identical homology. Amino acids are represented by the 1-letter code. X, is undetermined.

**Figure S2.** TRBJ gene segment homology. Alignment of the nucleic acid sequences of the human, macfas and macmul TRBJ genes. Dots represent identical homology.

**Figure S3.** Single cell analysis of lung mononuclear cells from Cynomolgus macaques. (A) TRAV and (B) TRBV repertoire of T cells from bronchoalveolar lavage fluid, before or after infection (BAL (PRE), BAL (POST)), uninvolved or involved lung tissue (UI and granuloma, respectively), or single cell suspensions (SC). The mean value of the average percentage of the UMI counts from each sample was plotted. Each point represents a different sample. *, expression not analyzed.

**Table S1**: Macfas TRAV functional status, nucleotide sequence and the translated sequence

**Table S2.** Macfas TRAJ sequence.

**Table S3**: Macfas TRBV functional status, nucleotide sequence and the translated sequence.

**Table S4**: Macfas TRDV and TRDJ functional status, nucleotide sequence and the translated sequence.

**Table S5:** Macfas TRGV/TRGJ functional status, nucleotide sequence and the translated sequence.

