## Supplementary material for "Identification and characterization of the T cell receptor (TCR) repertoire of the Cynomolgus macaque (*Macaca Fascicularis*)": Supplemental Figures S1-3.pdf

Figure S1

```
TRAC
macfas 1   XIQNPDPAVYQLRGSKSNDTSVCLFTDFDSVMNVSQSKDSDVHITDKTVLDMRSMDFKSNGAVAWSNKSDFACTSAFKDS
macmul 1   ................................................................................
homsap 1   .............D...S.K..........QT..........Y.................S............AN..NN.

macfas 81  VIPADTFFPGTESVCDANLVEKSFETDMNLNFQNLSVIGFRILLLKVAGFNLLMTLRLWSS
macmul 81  .............................................................
homsap 81  I..E.....SP..S..VK.........T.................................

TRBC1
macfas 1   EDLKKVFPPKVAVFEPSEAEISHTQKATLVCLATGFYPDHVELSWWVNGKEVHSGVSTDPQPLKEQPALEDSRYCLSSRL
macmul 1   ................................................................................
homsap 1   ...N.....E..........................F................................N..........

macfas 81  RVSATFWHNPRNHFRCQVQFYGLSEDDEWTEDRDKPITQKISAEVWGRADCGFTSVSYQQGVLSATILYEILLGKATLYA
macmul 81  ................................................................................
homsap 81  .......Q.................N....Q..A..V..IV...A...................................

macfas 161 VLVSALMLMAMVKRKDF
macmul 161 .................
homsap 161 ......V..........

TRBC2
macfas 1   EDLKKVFPPKVAVFEPSEAEISHTQKATLVCLATGFYPDHVELSWWVNGKEVHSGVSTDPQPLKEQPALEDSRYCLSSRL
macmul 1   ...................................................................T............
homsap 1   ....N....E...........................................................N..........

macfas 81  RVSATFWHNPRNHFRCQVQFYGLSEDDEWTEDRDKPITQKISAEAWGRADCGFTSESYQQGVLSATILYEILLGKATLYA
macmu  81  ................................................................................
homsap 81  .......Q.................N....Q..A..V..IV.......................................

macfas 161 VLVSALVLMAMVKRKDS--
macmul 161 .................--
homsap 161 .................RG

TRGC2
Macfas 1   DKHLDADVSPKPTIFLPSIAETNLHKAGTYLCLLEKFFPDVIEIHWQEKNSNKVLKSQEGNTMKTNNTYMKFSWLTVPEK
Macmul 1   ................................................................................
Homsap 1   ..Q...................K.Q...............I.K......K..TI.G..........D............E

Macfas 81  SLDKEHRCIVRHENNRNGVDQEIIFPPIKT----------------DVTTVDPKDSFSKDANDALLLQLTNTSAYYMYLL
Macmul 81  ..............................----------------.D.......NY.E.....Q...............
Homsap 81  ...............K..I...........DVTTVDPKYNYSKDAN..I.M....NW......T................

Macfas 145 LLLKSEVYFAIIAVCLLRRTAVCCNGERS
Macmul 145 ..V..........................
Homsap 161 .....V......TC...G...F.....K.

TRDC
Macfas 1   -RQPHTKPSVFVMKNGTNVACLVKDFYPKDIRINLESSKKITEFDPAIVVSPSGKYNAVKLGQYADSNSVTCSVQHNKEV
Macmul 1   X...............................................................................
Homsap 1   XS......................E..........V.............I............K.E...........DNKT

Macfas 80  VYSTDFEVKTNSTDHLKPTETENTKQPSKSCHEPKAIVHAEKVNMMSLTVLGLRMLFAKSVAINFLLTAKLLFL*
Macmul 81  ...................................--......................................
Homsap 81  .H........D....V..K.............K......T...................T..V........F...
```

```
TRBJ1

1.1     macfas TGAACACTGAAGCTTTCTTTGGACAAGGCACCAGACTCACAGTTGTAG
        macmul ............................................T...
        homsap ................................................

1.2     macfas CTAACTATGACTACACCTTCGGTTCAGGGACCAAGTTAACTGTTGTAG
        macmul ................................................
        homsap .........G...............G.......G......C.......

1.3     macfas TTCTGGAAACACCGTGTATTTTGGAGAGGGAAGTCGGCTCACTGTTGTAG
        macmul ..................................................
        homsap C............A.A..................T...............

1.4     macfas CAACTAATGAAAAACTGTTTTTTGGCAGTGGAACCCAGCTCTCTGTCTTGG
        macmul ...................................................
        homsap ...................................................

1.5     macfas TAGCAATCAGCCCCAGTATTTTGGAGATGGCACTCGACTCTCCGTCCTAG
        macmul ..................................................
        homsap ................C.......T.....G............A......

1.6     macfas CTCCTATAATTCGCCCCTCCACTTTGGGAACGGGACCAGGCTCACTGTGACAG
        macmul ...................................................G.
        homsap ............A.................T......................

TRBJ2

2.1     macfas CTCCTACAATGAGCAGTTCTTTGGGCCAGGCACACGGCTCACCGTGCTAG
        macmul ..................................................
        homsap .....................C........G...................

2.2     macfas CCAACACCGCGCAGCTGTTCTTTGGAGAAGGCTCTAGGCTGACCGTGCTGG
        macmul .T.................................................
        homsap .G.......G.G.......T..........................A....

2.2P    macfas CTGAGAGGCGCTGCTGGGCGTCTGGGCCGAGGACTCCTGGTTCTGG
        macmul .................................G............
        homsap ...........................G..................

2.3     macfas AGCACAGATCCGCAGTATTTTGGCCCAGGCACCCGGCTGACAGTGCTCG
        macmul .................................................
        homsap .........A.......................................

2.4     macfas AGCC-AAAACACTCAGTACTTCGGCGCCGGGACCCGGCTCTCAGTGCTGG
        macmul ....-.............................................
        homsap ....A......T......................................

2.5     macfas ACCAAGAGACCCAGTACTTCGGACCAGGCACGCGGCTCCTGGTGCTCG
        macmul ................................................
        homsap ......................G.........................

2.6     macfas CTCTGGGGCCAGCGTCCTGACTTTCGGGGCCGGCAGCCGGCTGACCGTGCTGG
        macmul .....................................................
        homsap ...........A.........................A...............

2.7     macfas CTCCTACGAGCAGTACTTCGGGCCGGGCACCAGGCTCACAGTCATAG
        macmul ...............................................
        homsap .......................................G....C..
```

Supplemental Figure S2

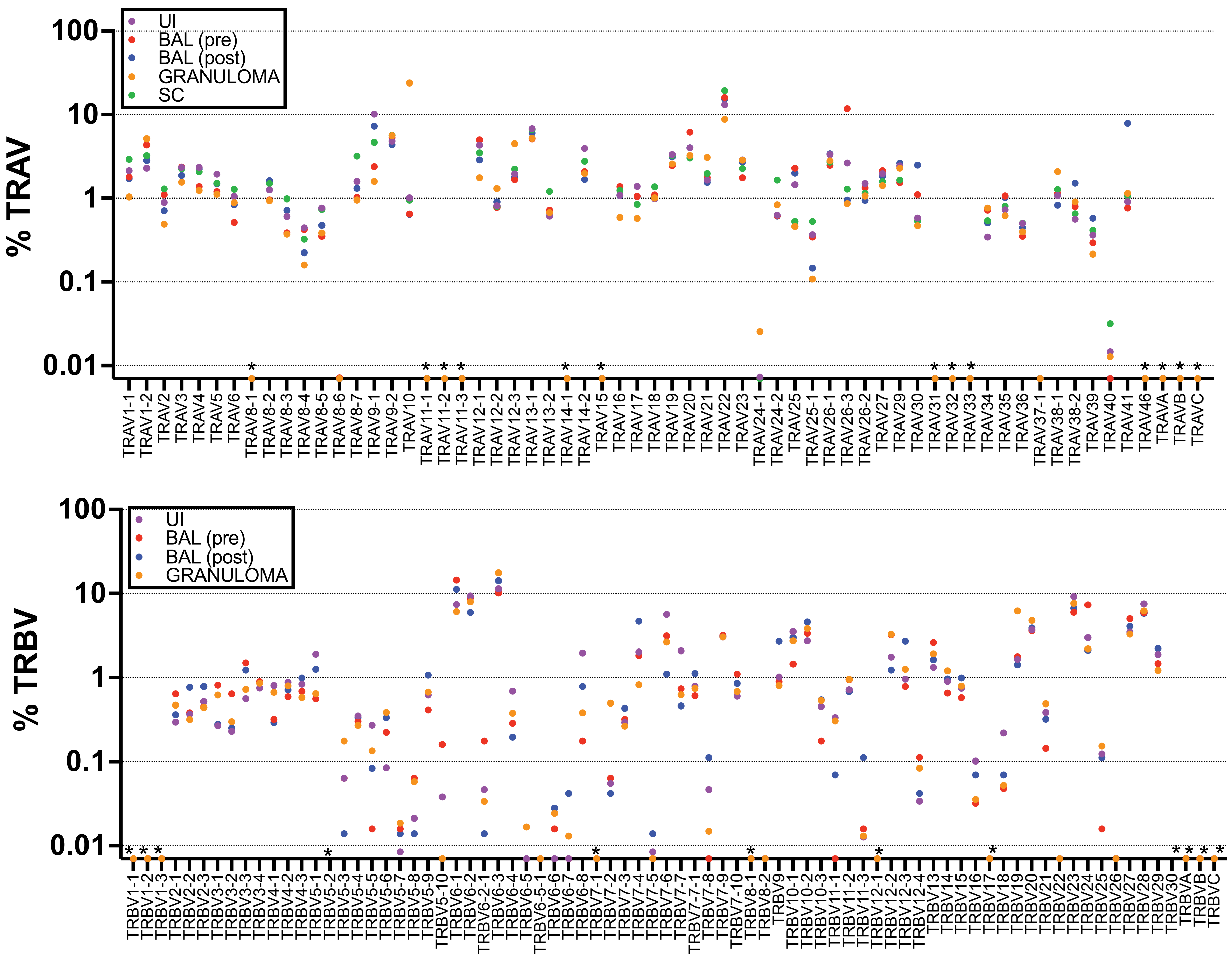


Supplemental Figure 3
